# Shared Graft Versus Leukemia Minor Histocompatibility Antigens in DISCOVeRY-BMT

**DOI:** 10.1101/2022.08.12.503667

**Authors:** Kelly S Olsen, Othmane Jadi, Sarah Dexheimer, Dante Bortone, Steven Vensko, Sarah Bennett, Hancong Tang, Marisa Diiorio, Tanvi Saran, David Dingfelder, Qianqian Zhu, Yiwen Wang, Christopher A Haiman, Loreall Pooler, Xin Sheng, Amy Webb, Marcelo C Pasquini, Philip L McCarthy, Stephen R Spellman, Eric Weimer, Theresa Hahn, Lara Sucheston-Campbell, Paul Armistead, Benjamin Vincent

## Abstract

T cell responses to minor histocompatibility antigens (mHAs) mediate graft versus leukemia (GvL) effects and graft versus host disease (GvHD) in allogeneic hematopoietic cell transplant (alloHCT). Therapies that boost T cell responses improve the efficacy of alloHCT; however, these have been limited by concurrent increases in the incidence and severity of GvHD. mHAs with expression restricted to hematopoietic tissue (GvL mHAs) are attractive targets for driving GvL without causing GvHD. Prior work to identify mHAs has focused on a small set of mHAs or population-level SNP association studies. We report here the discovery of a large set of novel GvL mHAs based on predicted peptide immunogenicity, restriction of expression to hematopoietic tissue or GvHD target organs, and degree of sharing among donor-recipient pairs (DRPs) in the DISCOVeRY-BMT dataset of 3231 alloHCT DRPs. The total number of predicted mHAs and count within each class of predicted mHAs significantly differed by recipient genomic ancestry group, with European American>Hispanic>African American for each. The number of mHAs also differed markedly by HLA allele, even among alleles of the same gene. From the pool of predicted mHAs, we identified the smallest sets of GvL mHAs needed to cover 100% of DRPs with a given HLA allele. We then used mass spectrometry to search for high population frequency mHAs for three common HLA alleles. We validated a total of 24 novel predicted GvL mHAs that cumulatively are found within 98.8%, 60.7%, and 78.9% of DRPs within DISCOVeRY-BMT that express HLA-A*02:01, HLA-B*35:01, and HLA-C*07:02 respectively. We also confirmed *in vivo* immunogenicity of one example novel mHA via coculture of healthy human CD8 T cells with mHA-pulsed dendritic cells. This work demonstrates that identification of shared mHAs is a feasible and promising technique for expanding mHA-targeting immunotherapeutics to larger numbers of patients.

## Introduction

Minor histocompatibility antigens (mHAs) are peptides derived from single nucleotide polymorphisms (SNPs) that differ between a tissue transplant recipient and donor, such that the mHA variant is expressed by the recipient only and presented on recipient major histocompatibility (MHC) molecules^1–4^. T cells that target these antigens are important mediators of the beneficial graft versus leukemia (GvL) effect and harmful graft versus host disease (GvHD) after allogeneic hematopoietic cell transplantation (alloHCT)^5–7^. AlloHCT is a standard of care for eligible patients with high risk Acute Myeloid Leukemia (AML), the deadliest form of leukemia in the US^8,9^. It is a highly effective treatment for AML in first complete remission (CR1) and reduces relapse risk by over 60% versus chemotherapy alone^8,10–12^. However, prognosis is poor for patients who relapse after alloHCT. Since the development of alloHCT, transplant clinicians have experienced the “transplanter’s dilemma”: interventions seeking to boost antileukemia T cell responses (i.e. increasing GvL) have also increased GvHD incidence, and interventions preventing GvHD have increased relapse rates^5,13–24^. A foundational problem in transplant immunology is separating GvL from GvHD in alloHCT recipients. One approach is to utilize GvL mHA-directed immunotherapies^25^. GvL mHAs are defined as mHAs that are only expressed in hematopoietic tissue, so T cell responses against them will lead to GvL effects without GvHD. Approximately 55 mHAs have been reported to date, including 13 validated GvL mHAs, and clinical trials of mHA-targeted cellular immunotherapies and vaccines have been performed^25–33^.

The majority of mHA discovery focuses on identifying personalized mHAs for individual transplant donor-recipient pairs (DRPs). The general workflow for these studies consists of isolation of activated T cells in a post-transplant recipient sample, T cell expansion and immunoprecipitation of their HLA-presented antigens, mass spectrometry (MS) to identify presented peptides, genome searching for a match for the peptide, and assessment of whether the source gene expression is restricted to hematopoietic tissue^9,34–38^. This approach is extremely laborious and identifies mHAs that may be only applicable for an individual DRP or a small number of patients. Thus, we report here an innovative approach that identifies GvL mHAs that are shared across many DRPs within the population.

## Methods

### Computational Methods

#### Study population

DRP sequencing and clinical data were derived from the DISCOVeRY-BMT (Determining the Influence of Susceptibility Conveying Variants Related to one-Year mortality after BMT) study, reported to CIBMTR from 151 transplant centers within the US^39–43^. Patients included in this study were treated for Acute Myeloid Leukemia (AML), Acute Lymphocytic Leukemia (ALL), and Myelodysplastic Syndrome (MDS) with alloHCT. Cohort 1 of DISCOVeRY-BMT consists of 2609 10/10 HLA-matched donor-recipient pairs (DRPs) treated from 2000-2008, while Cohort 2 consists of 572 10/10 HLA-matched DRPs treated from 2009-2011 and 351 8/8, but <10/10, HLA-matched DRPs treated from 2000-2011^42^. DRPs were excluded if the grafts were cord blood grafts or T cell-depleted, or SNP data was not available for both donor and recipient. For antigen prediction analyses, all patients were combined. Applicability of some predicted mHAs may vary by disease type due to expression levels of the source genes in individual patients or diseases.

All patients included in the DISCOVeRY-BMT study provided informed consent to be included in the Center for International Blood and Marrow Transplant Research (CIBMTR) registry. Genotyping was performed as previously described using the Illumina HumanOmni Express chip^42–44^. SNP quality control was performed and variants with minor allele frequency (MAF) < 0.005 were removed, leaving 637,655 and 632,823 measured SNPs for Cohort 1 and Cohort 2 respectively^40^. We calculated genetic distance between each DRP based on SNP array data^45^. Genomic ancestry was calculated via principal component analysis. Principal components were constructed using a set of independent SNPs in all patients self-declaring White, European, or Caucasian race and Non-Hispanic ethnicity. Mean values for the first three eigenvectors were determined and individuals with any of the first three eigenvectors greater than two standard deviations from each mean value were excluded. This was repeated for individuals self-declaring Black or African race and Non-Hispanic ethnicity, and for individuals declaring Hispanic ethnicity^43,46^. For this work, three genomic ancestry groups were assessed, including EA, Hispanic (HIS), and African American (AA), due to the small numbers of self-reported Asian American and Native American patients in the dataset. Patients that self-reported as Asian American and Native American were included in mHA prediction work but genomic ancestry was not calculated and these patients were excluded from ethnicity analyses. Student’s T tests and Chi-squared tests were performed to assess differences in number of predicted mHAs between groups.

#### Minor histocompatibility antigen prediction

Minor mismatches were defined as SNP loci where the graft recipient allele and the donor differed, and mHAs were the predicted peptides from the recipient allele of the minor mismatch^1^. All possible peptides of lengths between 8 and 11 amino acids resulting from SNP mismatches within every DRP were screened for predicted binding affinity against the recipient HLA class I alleles and expression of the source gene. We filtered for peptides with a peptide/HLA dissociation constant <500nM using NetMHCpan and >1 transcripts per million (TPM) in AML RNAseq data obtained from The Cancer Genome Atlas^47^. Peptides were called mHAs if they fit these criteria. Where multiple length peptides derived from the same SNP met the filter, these cases were reduced to the shortest version of the peptide. This avoids counting peptides with identical core sequences as separate entities, as patients containing a specific SNP will likely have each length version of that SNP-derived peptide. mHAs were then subcategorized based on source protein mRNA expression level. Peptides were labeled “GvL” if they showed expression levels of >50TPM in AML cells and <50TPM in GvHD target organs including skin, liver, and colon. “GvH” label indicates levels of <50TPM in AML cells and >50TPM in GvHD target organs. “Both” denotes >50TPM in both AML cells and GvHD target organs. Peptides with a “GvL” label were considered for further analysis, while peptides with tags of “Both” and “GvH” were excluded. This resulted in 1,867,836 potential mHAs of interest across 3231 DRPs.

#### Minimal set coverage calculation

We developed a greedy algorithm to resolve the maximum set coverage problem and generate ranked lists of the most commonly shared mHAs for recipients with each of our HLA alleles of interest. This algorithm generates a list of the minimal set of peptides such that every DRP with a given HLA allele in the DISCOVeRY-BMT dataset contains at least one of these mHAs. In short, the algorithm ranks every peptide within a given HLA by the study population frequency in descending order. The peptide with the highest frequency is selected and added to the mHA set, then population frequency of every peptide is recalculated using only DRPs that do not contain an mHA in the set and the new highest frequency peptide is selected. This process is repeated until 100% of DRPs are represented by an mHA in the set. For MS validation, we selected the peptides from the set that have non-zero RNAseq coverage of the gene that contain them in the cell line being utilized for validation. We added additional peptides for analysis by filtering the peptides not selected by the greedy algorithm for non-zero expression of the source gene, ranking in descending order of noncumulative population coverage by each peptide, then selecting the necessary number of peptides to bring the total list for mass spectrometric (MS) validation to 40 peptides as this was a feasible search size for MS.

Three HLA alleles were selected for mHA MS validation based on high frequency in US ethnic groups and to include representative alleles for HLA A, HLA B, and HLA C. HLA-A*02:01 is the most common HLA-A allele among Caucasians, African Americans, and Hispanics within the United States and third most common among Asians and Pacific Islanders, and is found within 28.4% of the total population of the United States. HLA-B*35:01 is the most common HLA-B allele among Asians and Pacific Islanders, is third most common among African Americans, and is fifth most common among Caucasians and Hispanics. It is found within 6.7% of the population of the United States. HLA-C*07:02 is the most common HLA-C allele among Hispanics within the United States, is second most common among Caucasians and Asians and Pacific Islanders, and is seventh most common among African Americans. It is found within 15.4% of the United States population^48^. One list of 40 peptides was searched for HLA allele B*35:01, and one list of 40 peptides was searched for HLA-C*07:02. Two samples were sent for MS for HLA-A*02:01. A 40 peptide search list was updated for the second sample to utilize updated cell line RNAseq data. In total, 67 peptides were searched for HLA-A*02:01.

### Experimental Methods

#### Cell lines

The AML cell lines used for MS were U937A2, the U937 cell line stably transfected to express HLA-A*02:01, NB4, which expresses HLA-B*35:01, and MONOMAC1, which expresses HLA-C*07:02^48^. Cell line HLA expression data was downloaded from the TRON cell lines portal and validated by the Clinical HLA Typing Laboratory at the University of North Carolina at Chapel Hill Hospitals, with some differences as reported (**Supplemental Figure 1**)^48,49^. Where discrepancies were found between the HLA haplotype on TRON and the clinical typing, the clinical typing result was used. Cell lines were maintained in culture with RPMI 1640 +10% FBS +1% penicillin-streptomycin +1% L-glutamine^50^.

#### RNAseq

RNA was isolated from 5 million cells per cell line using the Qiagen AllPrep DNA/RNA extraction kits (Qiagen, 80004). Nucleic acid quantification and quality detection were performed and RNA sequencing libraries were generated via Illumina Stranded mRNA prep. Paired-end sequencing was run on a NovaSeq SP 2×150 with the NovaSeq 6000 SP Reagent Kit v1.5 (NovaSeq, 20028400) and aligned to reference genome GRCh37, as SNP locations were called using this reference genome. Differential gene analysis was performed (**Supplemental Figure 2**).

#### Immunoprecipitation and Mass Spectrometry

Cell lines were expanded to 1-5×10^8^ per sample. Cells were centrifuged and washed with PBS twice, followed by treatment with 1x cOmplete Mini EDTA-free Protease Inhibitor Cocktail tables prepared in PBS (Roche, 11836170001). Cells were centrifuged and supernatant removed, and cell pellets snap frozen in liquid nitrogen and placed at −80C. Frozen pellets were sent to Complete Omics Inc for immunoprecipitation and antigen validation and quantification by mass spectrometry through Valid-Neo® platform^51^. Pellets were processed into single-cell frozen powder and then lysed. Peptide-HLA complexes were immunoprecipitated using Valid-NEO® neoantigen enrichment column pre-loaded with anti-human HLA-A, B, C antibody clone W6/32 (Bio-X-Cell). After elution, dissociation, filtration and clean up, peptides were lyophilized before further analysis. Transition parameters for each epitope peptide were examined and curated through Valid-NEO® method builder, an AI-based biostatistical pipeline. Ions with excessive noise due to co-elution with impurities were further optimized or removed. To boost detectability, a series of computational recursive optimizations of significant ions were conducted. Each neoantigen sequence was individually detected and quantified in a high-throughput manner through a Valid-NEO® modified mass spectrometer.

#### mHA Immunogenicity Assessment

Human donor Leukopaks were obtained (Gulf Coast Regional Blood Center) and genotyped for HLA*A:02 via flow cytometry with purified anti-human HLA-A2 antibody clone BB7.2 (BioLegend). HLA-A*02 positive samples were selected and dendritic cells were generated via plate adherence and pulsed with mHAs that are not endogenous to the sample (Peptide 2.0 Inc). Naïve CD8 T cells were isolated and cocultures were initiated with mHA-pulsed DCs and naïve CD8 T cells at a 1:4 ratio and maintained for two weeks in culture in RPMI + 10% Human Serum +1% penicillin-streptomycin +1% L-glutamine. Presence of mHA-specific T cells was assessed via flow cytometry with mHA tetramer staining. Tetramers were generated with Flex-T™ HLA-A*02:01 Monomer UVX (BioLegend, 280004) and fluorophore-conjugated streptavidin. Cells were also stained with the following: FVS700 live/dead (BD Biosciences), CD8-BV421 (BioLegend, Clone: SK1). Cells cultured with immunodominant Influenza A Virus M1_58-66_ HLA-A*02:01-binding influenza peptide and stained with Flu-M1_58-66_ tetramer (designated as Flu) were utilized as a positive control, and cells stained with tetramer exposed to UV light with no peptide (UV only) were utilized as negative control^52^. Gating strategy is shown (**Supplemental Figure 4**).

## Results

### Patient Characteristics and mHA predictions

Characteristics of patients in DISCOVeRY-BMT are shown in **Supplemental Table 1**^41^. 60% of DISCOVeRY-BMT patients had a diagnosis of AML, while the remainder had diagnoses of acute lymphoblastic leukemia (ALL) or myelodysplastic syndrome (MDS). Age distribution in DISCOVeRY-BMT reflects the age distribution of the predominant disease type in this dataset, AML, with 60% of alloHCT recipients older than 40 years of age. Most transplant recipients in the dataset received bone marrow-derived grafts, while some received peripheral blood stem cell grafts. Using the SNP typing data from these patients, we predicted a total of 9,241,788 mHAs in the DISCOVeRY-BMT dataset.

The number of predicted mHAs did not vary by disease type (**Figure 1A**). The self-reported ethnicity and genomic ancestry of alloHCT recipients in this dataset mirror the general distribution of alloHCT recipients in the US, with a predominance of patients with European American (EA) ancestry^53^.

**Figure 1:**
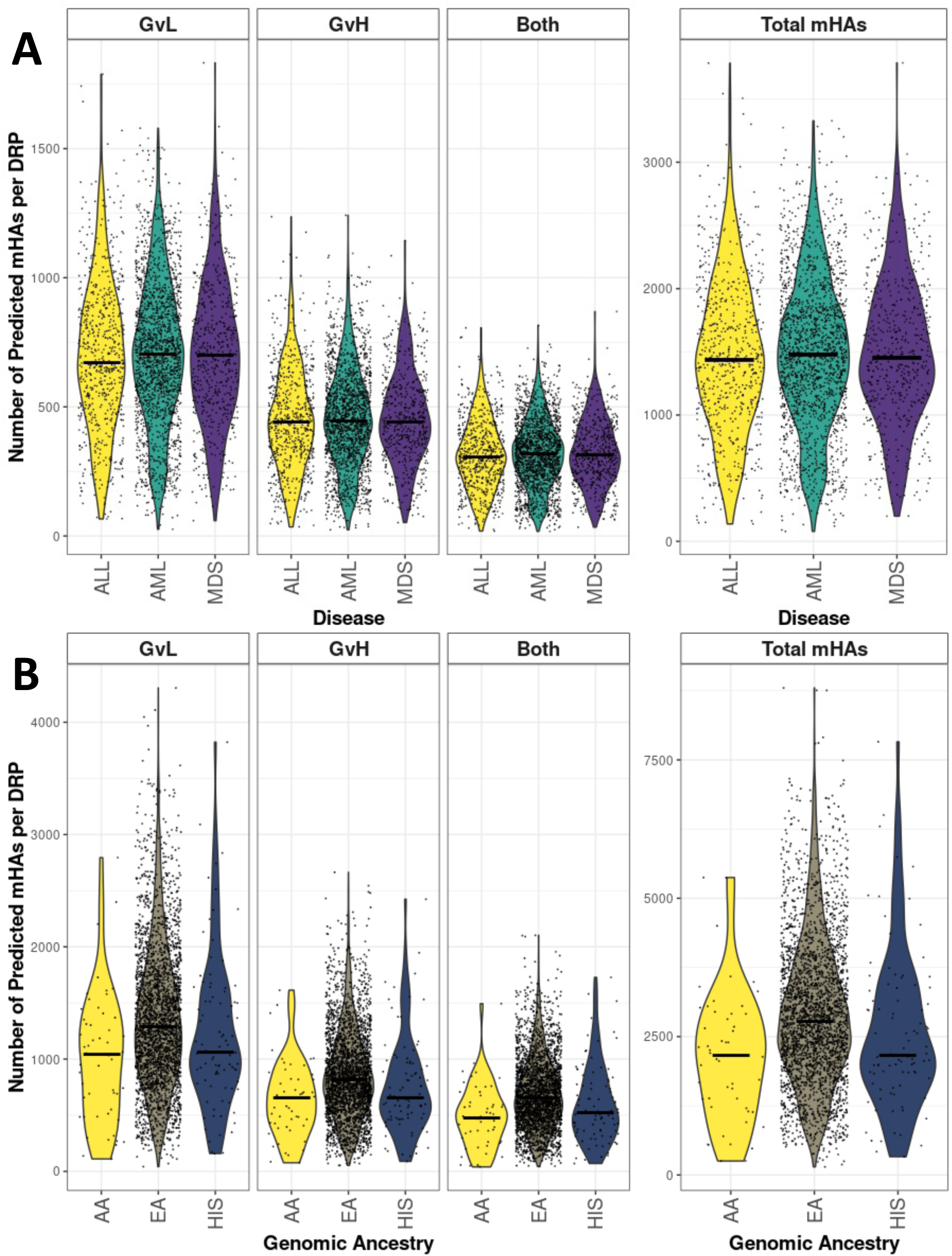
Predicted mHAs by disease type and genomic ancestry. 1a) shows number of each category of predicted mHA per donor-recipient pair (DRP) by disease type. “GvL” denotes expression in leukemia cells, “GvL” denotes expression in GvH target organs, and “both” denotes expression in both. 1b) shows number of each category of predicted mHA per DRP by genomic ancestry, including only patients who did not self-report as Asian American or Native American.

A large number of mHAs were predicted for each genomic ancestry group assessed in this study, with 75918 total predicted mHAs for EA, 27557 mHAs for AA, and 39272 mHAs for HIS (**Figure 1B**). mHAs were then assigned tags based on expression of the source gene in AML and GvHD target tissues. The mean total predicted mHAs per DRP across all ethnicities was 1476, with a mean of 704 predicted GvL mHAs] Number of predicted mHAs significantly differed by genomic ancestry group, with EA>HIS>AA for number mHAs labeled as GvL, GvH, and both as well as total mHAs per DRP.

### Predicted GvL mHAs were identified for 56 HLA alleles found in DISCOVeRY-BMT alloHCT recipients

A total of 23 HLA A alleles, 26 HLA B alleles, and seven HLA C alleles were represented in DISCOVeRY-BMT. The total number of predicted mHAs that bind each allele varied widely: from 82 to 11017 for HLA-A alleles, 19 to 8585 for HLA-B alleles, and 946 to 7537 for HLA-C alleles (**Figure 2A/B/C**). However, our method predicted GvL mHAs for every HLA allele represented within DISCOVeRY-BMT, with 35 GvL mHAs predicted for HLA-A*26:03, 10 for HLA-B*46:01, and 483 for C*06:02, the A, B, and C alleles with the fewest predicted GvL mHAs. Next, we looked at the proportion of mHAs classified as GvL, GvH, or both for each HLA allele. GvL mHA comprised approximately half of all predicted mHAs for each HLA allele (**Figure 2D/E/F**).

**Figure 2:**
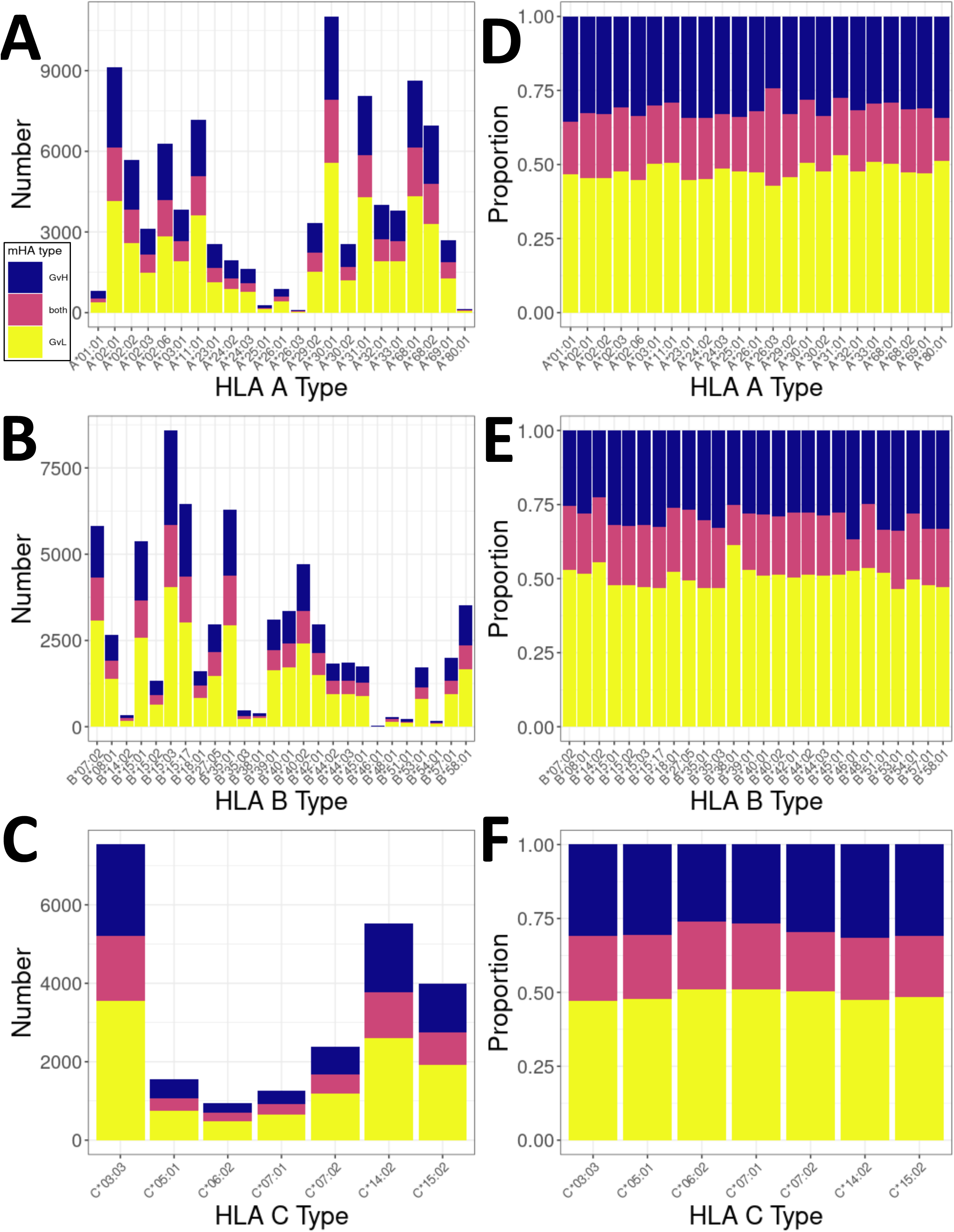
Number and proportion of predicted mHAs by HLA allele within study population. mHAs classed as “GvL” broadly represent mHAs that are desirable to target for anti-leukemia effects with minimal GvHD. mHAs classed as “GvH” represent mHAs that are undesirable to target as they are predicted to correspond to GvHD and no GvL effects. “Both” category represents peptides that are predicted to lead to both GvL and GvH effects. 2a) shows counts of each predicted class of mHA for HLA A alleles represented in patient dataset. 2b) shows counts for HLA B alleles represented in patient dataset. 2c) shows counts for HLA C alleles represented in patient dataset. 2d) shows proportion of predicted mHAs corresponding to each mHA class for HLA A alleles. 2e) shows proportion for HLA B alleles. 2f) shows proportion of HLA C alleles.

### Genetic distance between donor and recipient does not correlate with number of predicted GvL mHAs

We next assessed whether overall genetic distance between donor and recipient correlates with predicted total mHAs or GvL mHAs. We saw a strong positive correlation between total number of mHA-encoding SNPs and number of predicted GvL mHAs (**Figure 3A**). We observed a narrow range of pairwise genetic distance across all DRPs within DISCOVeRY-BMT (**Figure 3B**), likely resulting from flattening of diversity due to the large number of rare SNPs genotyped. Still, distance values were consistent with previously reported data for healthy pairs^45^. We observed no correlation between genetic distance and predicted GvL mHAs (**Figure 3C**) or total mHAs (**Figure 3D**).

**Figure 3:**
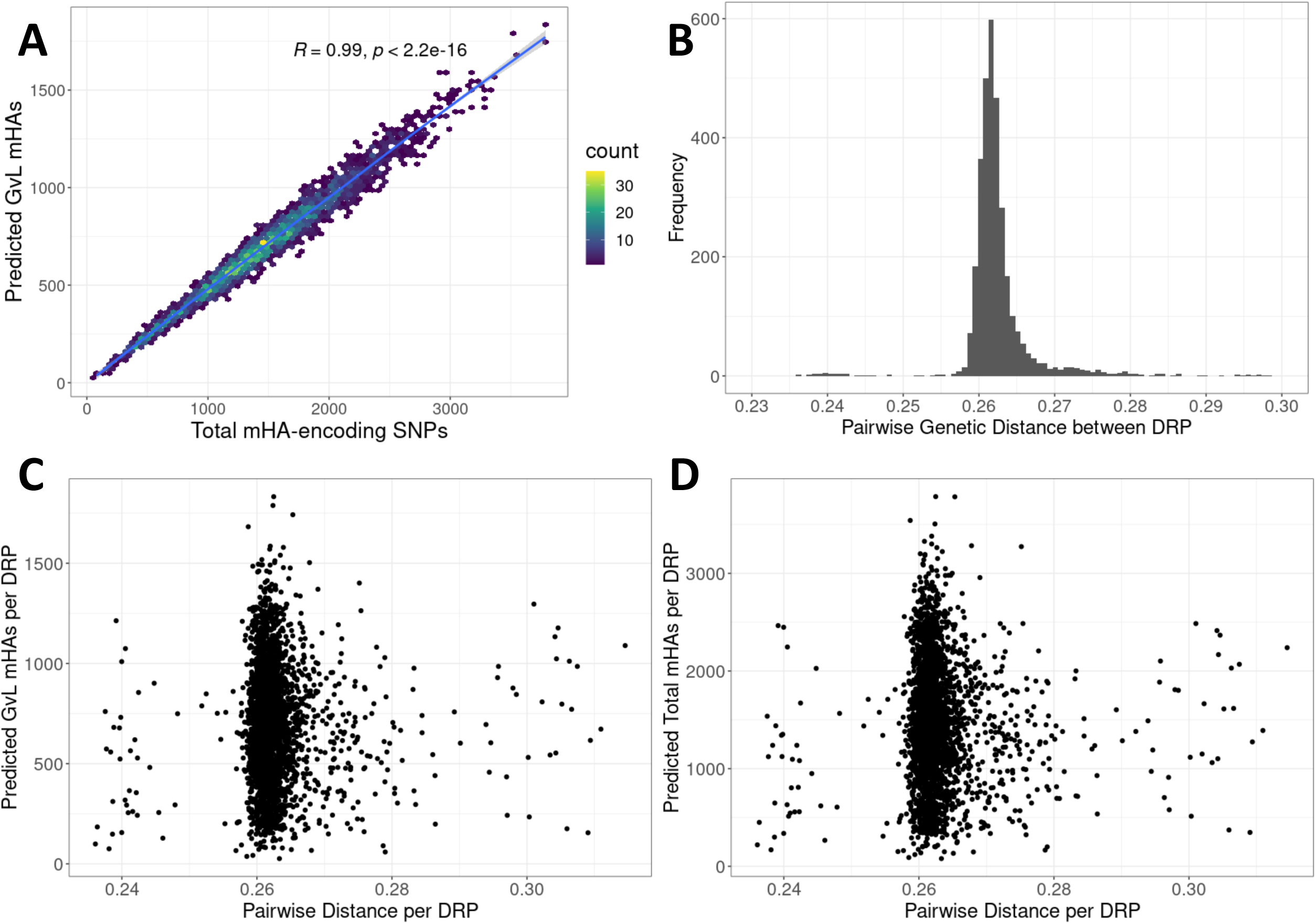
Degree of genetic distance versus number of predicted GvL mHAs by DRP in DISCOVeRY-BMT dataset. 3a) shows number of total SNPs that differ and are predicted to lead to an mHA versus number of predicted GvL mHAs per patient. 3b) shows distribution of pairwise distance values for every DRP in the DISCOVeRY-BMT dataset. Pairwise genetic distance value is calculated as the mean of (1-.5(number of shared alleles at SNP locus)) for every genotyped SNP locus for a DRP. 3c) shows pairwise genetic distance versus number of predicted total mHAs per DRP. 2d) shows pairwise genetic distance versus number of predicted GvL mHAs per DRP.

### Most predicted mHAs are private, but a small number are widely shared among patients with any given HLA allele

We evaluated sharing of predicted mHAs within the DISCOVeRY-BMT cohort. Of our predicted mHAs, the majority were found within a few recipients. However, 38.7% of our predicted mHAs were shared by 1% or more of the study population, and 4% were shared by 10% or more of the study population (**Figure 4A**). Next, we assessed sharing of mHAs within individual HLA alleles. For the three HLA alleles focused on in this work, the population frequency of predicted mHAs shows a bimodal distribution. Most mHAs are unshared, but a group of mHAs covers approximately 20-30% of patients (**Figure 4B-D**). Finally, we assessed predicted mHA frequency across all HLA alleles represented by greater than 0.5% of DISCOVeRY-BMT patients. The same bimodal distribution of mHA population frequency was observed across most HLA alleles (**Figure 4E**).

**Figure 4.**
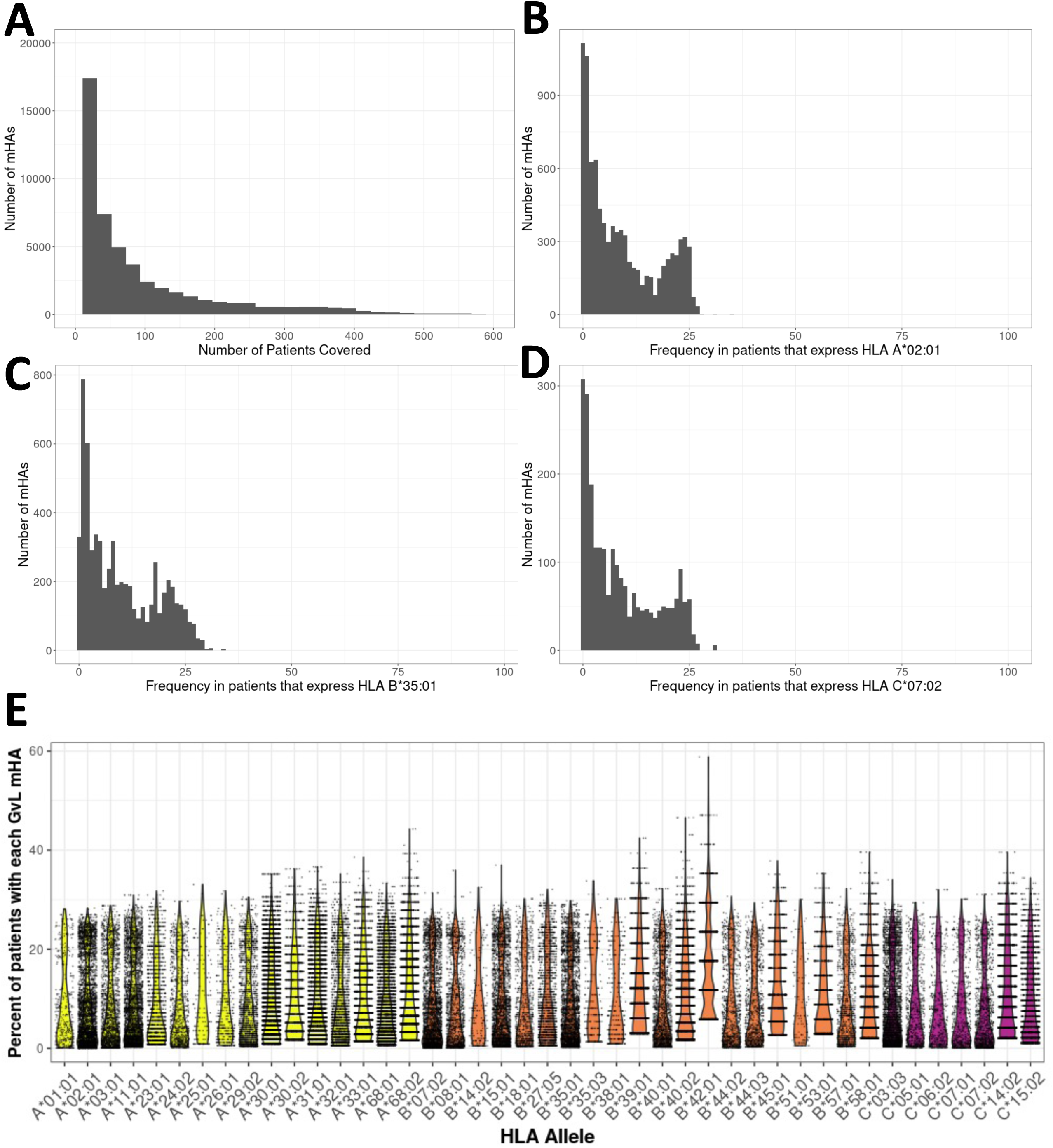
Degree of sharing of predicted mHAs across study population. 4a) shows distribution of predicted mHAs by the number of patients in the DISCOVeRY-BMT cohorts that possess them. 39.1% of predicted mHAs are unique to a single patient or shared by ten or fewer patients. However, 38.7% of predicted mHAs are shared by 32 patients or more, corresponding to 1% of the study population. 1602 predicted mHAs (4.0% of total GvL mHAs) are shared by 323 patients or more, corresponding to 10% of the study population. 4b) shows distribution of predicted HLA A*02:01 mHAs by population frequency in patients that express HLA A*02:01. 4c) shows distribution of predicted HLA B*35:01 mHAs by population frequency in patients that express HLA B*35:01. 4d) shows distribution of predicted HLA C*07:02 mHAs by population frequency in patients that express HLA C*07:02. 4e) shows percentage of DISCOVeRY-BMT cohort with each HLA allele covered by each predicted GvL mHA that binds that HLA allele, for all HLA alleles representing greater than 0.5% of DISCOVeRY-BMT patients.

### For three HLA alleles common in prevalent United States ethnic groups, 11-15 GvL mHA peptides cover 100% of patients in DISCOVeRY-BMT that express the given allele

By identifying a minimal set of GvL mHA peptides with cumulative 100% population coverage, we can discover the minimum number of GvL mHAs for a GvL mHA-directed therapy to apply to all patients with a specific HLA allele. By quantifying the size of these lists for the most common HLA alleles in genomic ancestry groups, we can estimate the total number of therapeutics needed to treat the majority of alloHCT recipients.

We selected three HLA alleles to generate minimal sets for, including HLA-A*02:01, HLA-B*35:01, and HLA-C*07:02. Together, these three HLA alleles represent a set of HLA alleles that is common within the US population and within the major ethnic groups found in the DISCOVeRY-BMT population. For the most common HLA allele in the US, HLA*A02:01, a set of fifteen GvL mHAs are needed to ensure that every DRP with this HLA allele has at least one of the fifteen (**Figure 5A**). The five peptides with the highest cumulative coverage are found within 80% of HLA*A02:01 DRPs, and only seven peptides are needed to reach 90% coverage. The non-cumulative population frequencies for each of these top fifteen peptides range from 19.4% to 28.3%. We obtained similar results with HLA-B*35:01: eleven peptides are needed to reach 100% population coverage and four peptides are needed to reach 80%, with non-cumulative population frequencies between 20.9% and 29.3% (**Figure 5B**). HLA*C07:02 also showed similar results, with fourteen peptides needed to reach 100% population coverage and five peptides needed to reach 80%. Noncumulative frequencies ranged from 19.3% to 31.1% (**Figure 5C**). A total of 40 peptides give 100% population coverage of three HLA alleles that are among the most common in four major ethnic groups in the United States.

**Figure 5.**
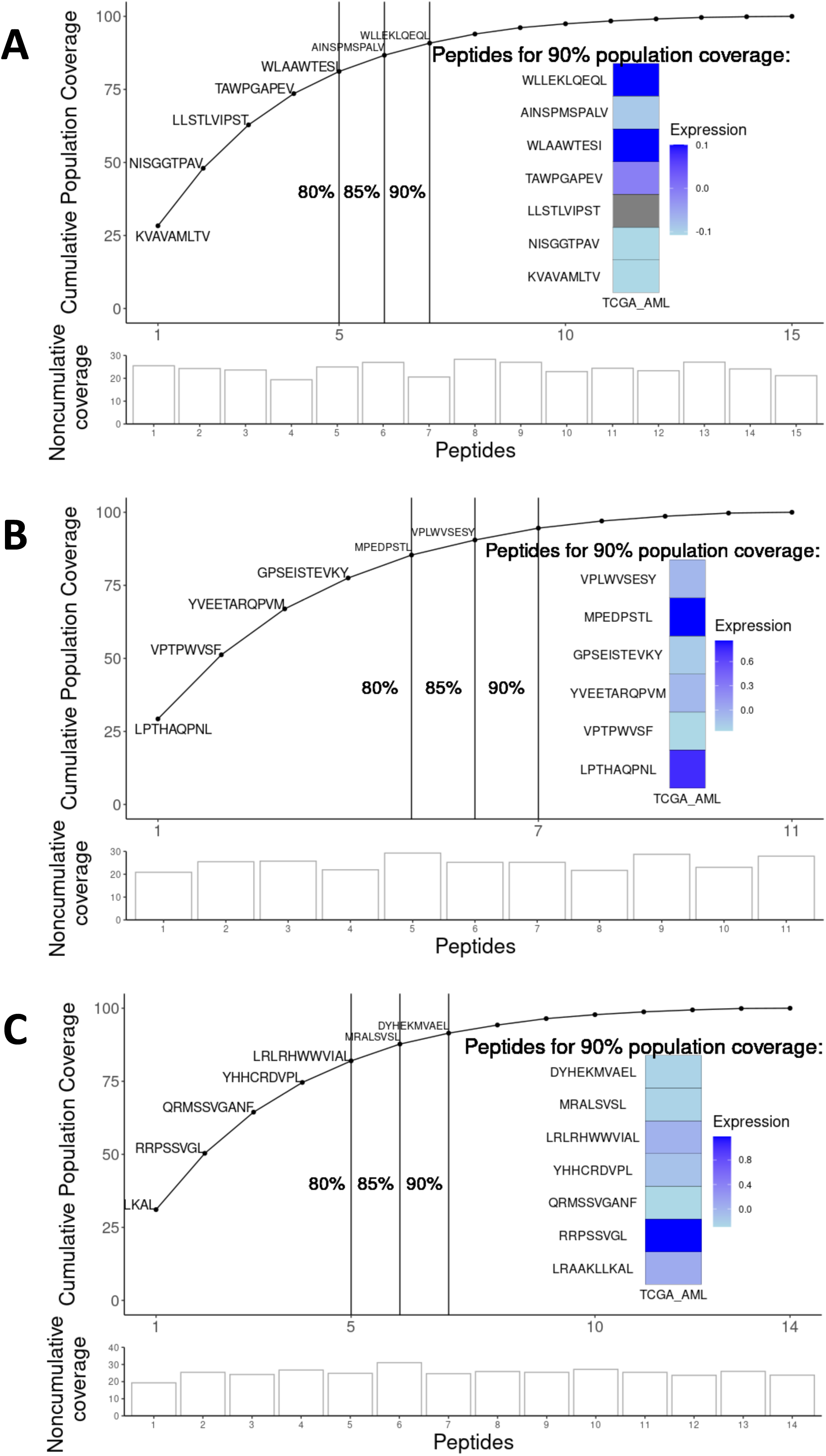
Patient population cumulative coverage by shared GvL mHAs. 5A) Coverage of DISCOVeRY-BMT patients with HLA*A02:01 allele with predicted GvL mHAs. Fifteen total GvL mHAs are needed to give 100% cumulative population coverage of patients with this HLA allele. Noncumulative independent population frequencies of each of the top fifteen peptides within the HLA*A02:01 population range from 19.4% to 28.3%. Inlaid heatmap shows z-scores of expression for the genes that contain each peptide from The Cancer Genome Atlas AML sample expression data (TCGA_AML). 5B) shows coverage of DISCOVeRY-BMT patients with HLA*B35:01 allele with predicted GvL mHAs. Eleven predicted GvL mHAs correspond to 100% cumulative population coverage for this allele. Noncumulative coverage by the top 11 peptides for this HLA allele range from 20.9% to 29.3%. 5C) shows coverage of DISCOVeRY-BMT patients with HLA*C07:02. Fourteen predicted GvL mHAs correspond to 100% cumulative population coverage for this allele. Noncumulative coverage for these mHAs range from 19.3% to 31.1%.

### 24 novel GvL mHAs were validated using mass spectrometry

We employed mass spectrometry to validate HLA presentation of predicted GvL mHAs. Of the 67 searched peptides for HLA-A*02:01 across two U937A2 cell line samples, we positively identified 17 peptides. Of the 40 searched for HLA-B*35:01 using an NB4 cell line we identified three peptides, and of the 40 searched for C*07:02 we identified five peptides. Representative spectra are shown for a heavy labeled peptide standard and endogenous identified peptide from an immunoprecipitated NB4 cell sample (**Figure 6A-B**). From the list of 17 validated peptides for HLA-A*02:01, peptide VLDIEQFSV is also known as UNC-GRK4-V and was previously identified by our group as a GvL mHA using the U937A2 cell line^1,29^. Mass spectrometry analysis was blinded to the peptide’s status as previously identified. As this peptide is previously known, a total of sixteen novel HLA-A*02:01 binding mHAs were discovered. These sixteen novel HLA-A*02:01-binding mHAs cumulatively cover 98.8% of HLA-A*02:01-positive patients in the DISCOVeRY-BMT dataset, with individual peptide population frequencies between 21.1% and 28.3% (**Figure 6C**). The three novel HLA-B*35:01 binding mHAs cover 60.7% of the HLA-B*35:01 positive DISCOVeRY-BMT population with population frequencies of 26.0-27.6% (**Figure 6D**). The five novel HLA-C*07:02 binding mHAs give cumulative HLA-C*07:02 positive DISCOVeRY-BMT patient coverage of 78.9%, with independent frequencies of 24.4-26.7% (**Figure 6E**). Characteristics of all novel mHAs are shown (**Table 1**). We also demonstrated immunogenicity of one example novel mHA, UNC-HEXDC-V, via tetramer staining of CD8 T cells cocultured with mHA-pulsed DCs (**Figure 6I**).

**Table 1.**
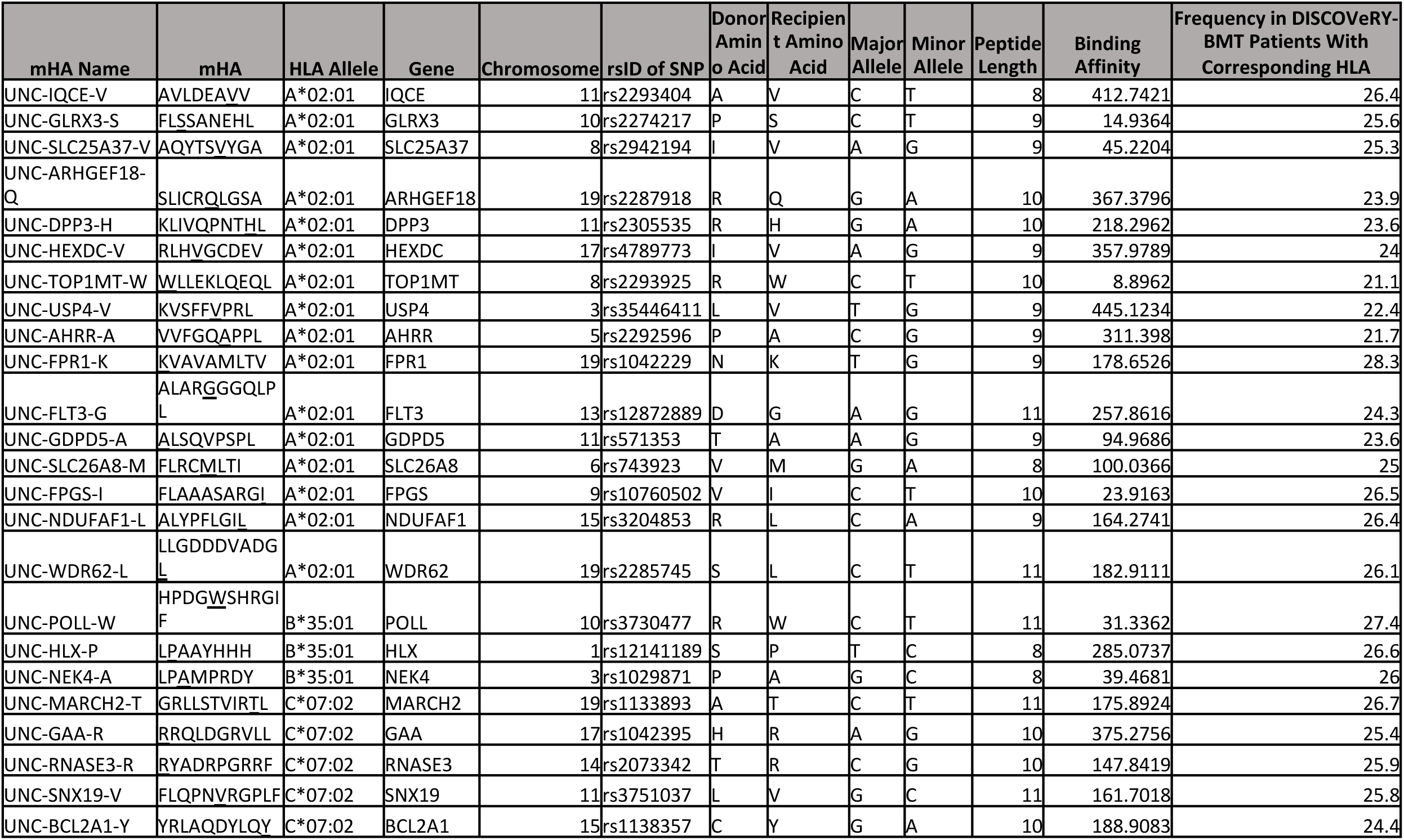
Novel mHA characteristics. 16 novel GvL mHAs that bind HLA A*02:01, three that bind B*35:01, and five that bind C*07:02 and were validated by mass spectrometry are shown.

**Figure 6.**
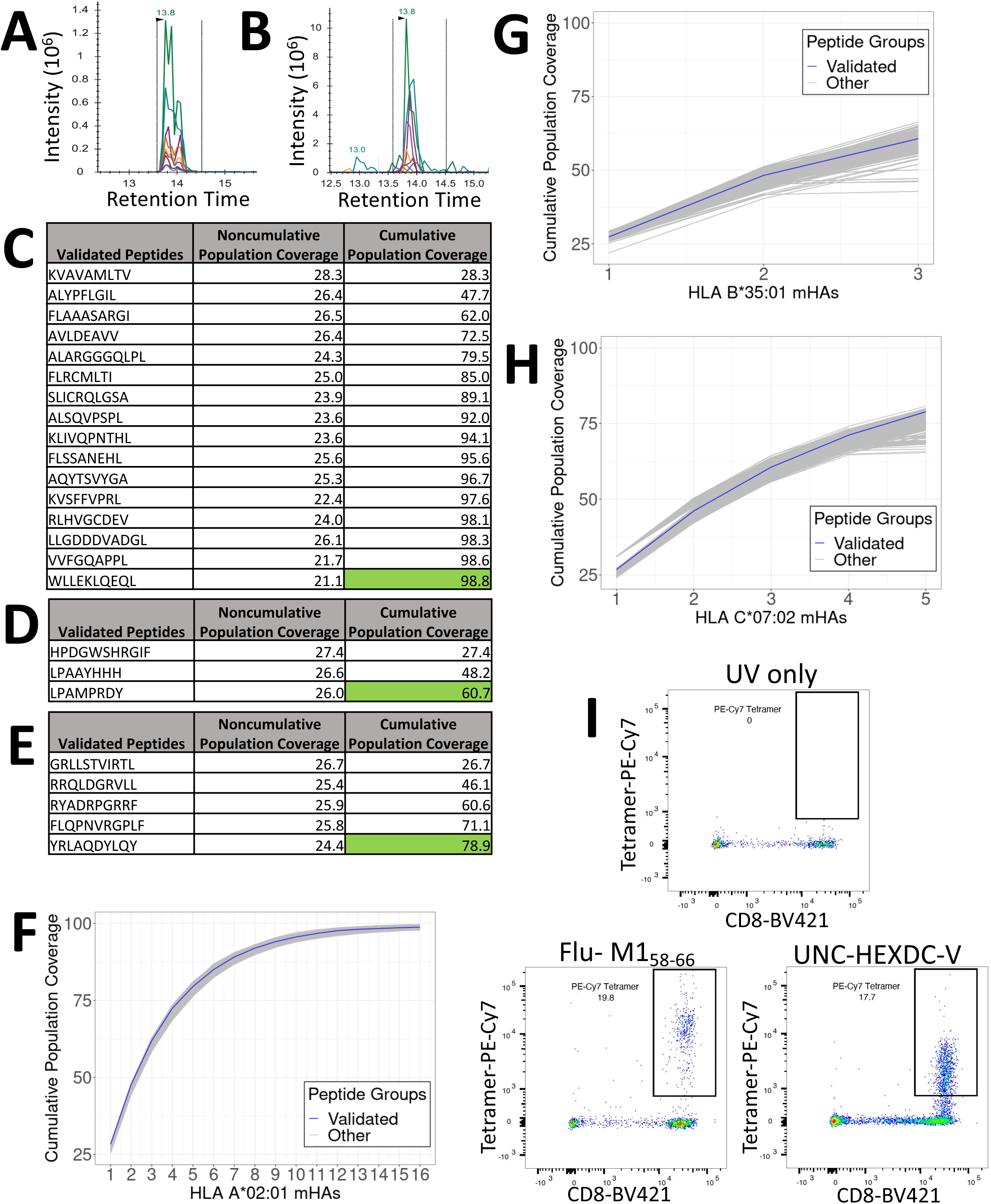
Mass spectrometry validation of predicted GvL mHAs for HLA A*02:01, B*35:01, and C*07:02. 6A) shows representative spectra for heavy-labeled peptide standard for HLA C*07:02-binding mHA LPAAYHHH. 6B) shows endogenous LPAAYHHH peptide identified from immunoprecipitated peptide sample from cell line MONOMAC1. 6C) shows all novel identified peptides from cell line U937A2 sample. “Noncumulative population coverage” shows percentage of DRPs expressing HLA A*02:01 within the DISCOVeRY-BMT dataset where the recipient expresses the mHA allele and donor does not. “Cumulative population coverage” shows output from greedy algorithm calculating total population coverage by each peptide and the ones preceding it, with a total of 98.8% population coverage by the ten peptides. 6D) shows all identified peptides from cell line NB4 sample, with 60.7% cumulative coverage of DRPs expressing HLA B*35:01 within the dataset by the three peptides. 6E) shows all identified peptides from cell line MONOMAC1, with 78.9% cumulative coverage of HLA C*07:02-expressing DRPs within the dataset. 6F) shows cumulative coverage by the 16 novel confirmed HLA A*02:01-binding mHAs and 1000 simulated sets of 16 peptides from the set of mHAs searched by mass spectrometry. Cumulative coverage by confirmed peptides is shown in blue, while each simulated run is shown as an individual gray line. 6G) shows cumulative coverage for the three confirmed HLA B*35:01-binding mHAs and 1000 simulated sets of three peptides. 6H) shows cumulative coverage for the five confirmed HLA C*07:02-binding mHAs and 1000 simulated sets of five peptides. 6I) shows flow cytometry staining from mHA immunogenicity experiment. “UV only” shows negative control stained with tetramer exposed to UV light with no peptide. “Flu-M15_8-66_” shows CD8 T cells cocultured with M1_58-66_ pulsed DCs, stained with M1_58-66_ tetramer. “UNC-HEXDC-V” shows CD8 T cells cocultured with novel mHA UNC-HEXDC-V stained with UNC-HEXDC-V tetramer.

To evaluate generalizability of our discovery process, we calculated the range of cumulative coverage that would be obtained with a subset of the number of peptides that we validated from the searched lists. For each HLA allele, 1000 random sets of peptides were selected from the searched peptide list and cumulative coverage by each set was calculated. The range of cumulative coverage by the 1000 random sets of 16 HLA-A*02:01 peptides was 97.4-99.7%, by the 1000 random sets of three HLA-B*35:01 peptides was 42.8-66.4%, and by the 1000 random sets of five HLA-C*07:02 peptides was 65.4-80.7% (**Figure 6F/G/H**).

## DISCUSSION

Discovery and characterization of novel mHAs may be crucial for enhancing immune monitoring in alloHCT, predicting clinical outcomes based on donor and recipient genetics, and improving outcomes by optimizing donor selection and/or specifically targeting GvL mHAs. We built upon previous work to perform the first population level survey of mHA peptides, taking a new approach by predicting mHAs common among recipients with diverse HLA alleles. This ensures that therapeutics targeting our newly identified mHAs would be applicable to as broad of a recipient population as possible. We evaluated mHAs for a total of 56 HLA A, B, and C alleles represented in 3,233 DISCOVeRY-BMT recipients. Despite large differences in total number of mHAs per HLA allele, approximately 50% of predicted mHAs for each HLA allele are GvL. Therefore, we expect that every HLA allele will present a set of GvL mHAs. The majority of GvL mHAs are shared among fewer than 10 patients in the dataset, highlighting the largely private nature of the mHA landscape. That said, for each HLA allele, we predicted a small number of highly shared mHA expressed by 20-25% of the recipient population. For all HLA alleles studied, 6-8 mHA peptides would cover >80% of recipients that express that allele, and 11-15 mHA would cover 100% of recipients. Conceptually, targeting a small number of shared GvL mHAs could treat a majority of alloHCT recipients regardless of race or ethnicity.

Using mass spectrometry, we validated a total of 24 novel GvL mHAs, an increase from the 13 GvL mHAs that have been discovered since Els Goulmy et al reported the first discovered GvL mHA, HA-1, in 1983^1,25,28,30–33^. The 16 novel GvL mHAs found for HLA-A*02:01 together cover 98.8% of HLA-A*02:01-positive patients in the DISCOVeRY-BMT dataset, the three for HLA-B*35:01 cover 60.7% of HLA-B*35-:01-positive DISCOVeRY-BMT patients, and the five for HLA-C*07:02 cover 78.9% of HLA-C*07:02-positive DISCOVeRY-BMT patients. Additionally, we confirmed immunogenicity of one predicted novel mHA, UNC-HEXDC-V, via tetramer staining of T cells cocultured with mHA-pulsed DCs. We expect that these novel mHAs will serve as future targets for antigen-directed therapeutics.

We genotyped seven DRPs from the UNC Lineberger Cancer Center Tissue Procurement Facility expressing HLA-A*02:01, one expressing HLA-B*35:01, and four expressing HLA-C*07:02 for the majority of the novel mHAs for the corresponding HLA alleles (**Supplemental Figure 3**). We found appropriate minor antigen mismatches for potential utilization of these mHAs in 58% of the genotyped DRPs, highlighting their utility for future work. This does not align perfectly with predicted coverage of DISCOVeRY-BMT patients with these mHAs, but this is likely explained by the small patient count and different patient population. However, most of the patients genotyped could utilize treatments targeting these mHAs. We also genotyped the seven HLA-A*02:01 positive DRPs for the previously known GvL mHAs HA-1 and UTA2-1 and discovered they were targetable in 0% of these DRPs. This emphasizes the expanded utility of finding shared mHAs over traditional methods.

Our study is limited in important ways. We biologically validated predicted GvL mHAs for three HLA alleles that were selected based on their high frequency of expression within diverse ethnic groups. In the future, mHAs for additional HLA alleles should be validated. Also, we validated GvL mHAs in a single AML cell line for each HLA allele. This is sufficient to establish that the mHAs are capable of being presented; however, antigen expression, HLA expression and antigen presentation efficiency will be heterogeneous across patient samples. Further studies of primary AML samples will be required to estimate the frequency of expression of each GvL mHA in AML. While this study includes *in vivo* validation of immunogenicity of one of our novel mHAs with a healthy donor sample, future work will identify mHA-specific T cells for all mHAs validated in this work. We will also assess T cell responses to the novel GvL mHAs in alloHCT recipients to better understand determinants of GvL mHA immunogenicity.

This work more than doubles the number of known validated GvL mHAs, and these mHAs are unique in being specifically identified for their high population prevalence in the corresponding HLA-expressing DRPs. Targeting these newly discovered mHAs could greatly expand the capacity for treatment of AML patients with GvL mHA-targeting immunotherapies.

## Acknowledgments

We want to acknowledge the participation of all the patients and donors who consented to the biorepository and research database, as well as all transplant centers which participated in the CIBMTR Database and Biorepository studies.

## References

1. Lansford JL, Dharmasiri U, Chai S, et al. Computational modeling and confirmation of leukemia-associated minor histocompatibility antigens. Blood Adv. 2018;2(16):2052–2062.

2. Griffioen M, van Bergen CAM, Falkenburg JHF. Autosomal Minor Histocompatibility Antigens: How Genetic Variants Create Diversity in Immune Targets. Front. Immunol. 2016;7:100.

3. Mullally A, Ritz J. Beyond HLA: the significance of genomic variation for allogeneic hematopoietic stem cell transplantation. Blood. 2006;109(4):1355–1362.

4. Bleakley M, Riddell SR. Exploiting T cells specific for human minor histocompatibility antigens for therapy of leukemia. Immunol. Cell Biol. 2011;89(3):396–407.

5. Horowitz MM, Gale RP, Sondel PM, et al. Graft-versus-leukemia reactions after bone marrow transplantation. Blood. 1990;75(3):555–562.

6. Gale RP, Horowitz MM, Ash RC, et al. Identical-Twin Bone Marrow Transplants for Leukemia. Ann. Intern. Med. 1994;120(8):646–652.

7. Martin PJ, Levine DM, Storer BE, et al. Genome-wide minor histocompatibility matching as related to the risk of graft-versus-host disease. Blood. 2017;129(6):791–798.

8. Loke J, Vyas H, Craddock C. Optimizing Transplant Approaches and Post-Transplant Strategies for Patients With Acute Myeloid Leukemia. Front. Oncol. 2021;11:666091.

9. Warren EH, Fujii N, Akatsuka Y, et al. Therapy of relapsed leukemia after allogeneic hematopoietic cell transplantation with T cells specific for minor histocompatibility antigens. Blood. 2010;115(19):3869–3878.

10. Cornelissen JJ, van Putten WLJ, Verdonck LF, et al. Results of a HOVON/SAKK donor versus no-donor analysis of myeloablative HLA-identical sibling stem cell transplantation in first remission acute myeloid leukemia in young and middle-aged adults: benefits for whom? Blood. 2007;109(9):3658–3666.

11. Schlenk RF, Döhner K, Krauter J, et al. Mutations and Treatment Outcome in Cytogenetically Normal Acute Myeloid Leukemia. N. Engl. J. Med. 2008;358(18):1909–1918.

12. Armistead PM, de Lima M, Pierce S, et al. Quantifying the survival benefit for allogeneic hematopoietic stem cell transplantation in relapsed acute myelogenous leukemia. Biol. Blood Marrow Transplant. J. Am. Soc. Blood Marrow Transplant. 2009;15(11):1431–1438.

13. Ferrara JLM, Levine JE, Reddy P, Holler E. Graft-versus-host disease. Lancet Lond. Engl. 2009;373(9674):1550–1561.

14. MacDonald KPA, Hill GR, Blazar BR. Chronic graft-versus-host disease: biological insights from preclinical and clinical studies. Blood. 2017;129(1):13–21.

15. Shlomchik WD. Graft-versus-host disease. Nat. Rev. Immunol. 2007;7(5):340–352.

16. Kosuri S, Herrera DA, Scordo M, et al. The Impact of Toxicities on First Year Outcomes after Ex-vivo CD34+ Selected Allogeneic Hematopoietic Cell Transplantation in Adults with Hematologic Malignancies. Biol. Blood Marrow Transplant. J. Am. Soc. Blood Marrow Transplant. 2017;23(11):2004–2011.

17. Scordo M, Shah GL, Kosuri S, et al. Effects of Late Toxicities on Outcomes in Long-Term Survivors of Ex-Vivo CD34+-Selected Allogeneic Hematopoietic Cell Transplantation. Biol. Blood Marrow Transplant. J. Am. Soc. Blood Marrow Transplant. 2018;24(1):133–141.

18. Urbano-Ispizua A, Carreras E, Marín P, et al. Allogeneic transplantation of CD34(+) selected cells from peripheral blood from human leukocyte antigen-identical siblings: detrimental effect of a high number of donor CD34(+) cells? Blood. 2001;98(8):2352–2357.

19. Anasetti C, Logan BR, Lee SJ, et al. Peripheral-blood stem cells versus bone marrow from unrelated donors. N. Engl. J. Med. 2012;367(16):1487–1496.

20. Lee SJ, Logan B, Westervelt P, et al. Comparison of Patient-Reported Outcomes in 5-Year Survivors Who Received Bone Marrow vs Peripheral Blood Unrelated Donor Transplantation: Long-term Follow-up of a Randomized Clinical Trial. JAMA Oncol. 2016;2(12):1583–1589.

21. Kröger N, Solano C, Wolschke C, et al. Antilymphocyte Globulin for Prevention of Chronic Graft-versus-Host Disease. N. Engl. J. Med. 2016;374(1):43–53.

22. Nash RA, Antin JH, Karanes C, et al. Phase 3 study comparing methotrexate and tacrolimus with methotrexate and cyclosporine for prophylaxis of acute graft-versus-host disease after marrow transplantation from unrelated donors. Blood. 2000;96(6):2062–2068.

23. Cutler C, Logan B, Nakamura R, et al. Tacrolimus/sirolimus vs tacrolimus/methotrexate as GVHD prophylaxis after matched, related donor allogeneic HCT. Blood. 2014;124(8):1372–1377.

24. Bejanyan N, Rogosheske J, DeFor TE, et al. Sirolimus and Mycophenolate Mofetil as Calcineurin Inhibitor-Free Graft-versus-Host Disease Prophylaxis for Reduced-Intensity Conditioning Umbilical Cord Blood Transplantation. Biol. Blood Marrow Transplant. J. Am. Soc. Blood Marrow Transplant. 2016;22(11):2025–2030.

25. Oostvogels R, Lokhorst HM, Mutis T. Minor histocompatibility Ags: identification strategies, clinical results and translational perspectives. Bone Marrow Transplant. 2016;51(2):163–171.

26. Meij P, Jedema I, van der Hoorn MAWG, et al. Generation and administration of HA-1-specific T-cell lines for the treatment of patients with relapsed leukemia after allogeneic stem cell transplantation: a pilot study. Haematologica. 2012;97(8):1205–1208.

27. Warren RL, Freeman JD, Zeng T, et al. Exhaustive T-cell repertoire sequencing of human peripheral blood samples reveals signatures of antigen selection and a directly measured repertoire size of at least 1 million clonotypes. Genome Res. 2011;21(5):790–797.

28. Goulmy E. Minor histocompatibility antigens: allo target molecules for tumor-specific immunotherapy. Cancer J. Sudbury Mass. 2004;10(1):1–7.

29. Dharamsiri U, Hunsucker SA, Vincent BG, et al. UNC-GRK4-1: An Allele Specific Cancer Testis Antigen Identified Through Genomic Screening. Blood. 2013;122(21):3246.

30. Amado-Azevedo J, Reinhard NR, van Bezu J, et al. The minor histocompatibility antigen 1 (HMHA1)/ArhGAP45 is a RacGAP and a novel regulator of endothelial integrity. Vascul. Pharmacol. 2018;101:38–47.

31. Choi EY, Choi K, Nam G, Kim W, Chung M. H60: A Unique Murine Hematopoietic Cell-Restricted Minor Histocompatibility Antigen for Graft-versus-Leukemia Effect. Front. Immunol. 2020;11:1163.

32. Kremer AN, Bausenwein J, Lurvink E, et al. Discovery and Differential Processing of HLA Class II-Restricted Minor Histocompatibility Antigen LB-PIP4K2A-1S and Its Allelic Variant by Asparagine Endopeptidase. Front. Immunol. 2020;11:381.

33. Pont MJ, van der Lee DI, van der Meijden ED, et al. Integrated Whole Genome and Transcriptome Analysis Identified a Therapeutic Minor Histocompatibility Antigen in a Splice Variant of ITGB2. Clin. Cancer Res. 2016;22(16):4185–4196.

34. van Bergen CAM, Kester MGD, Jedema I, et al. Multiple myeloma–reactive T cells recognize an activation-induced minor histocompatibility antigen encoded by the ATP-dependent interferon-responsive (ADIR) gene. Blood. 2007;109(9):4089–4096.

35. Brickner AG, Warren EH, Caldwell JA, et al. The Immunogenicity of a New Human Minor Histocompatibility Antigen Results from Differential Antigen Processing. J. Exp. Med. 2001;193(2):195–206.

36. Brickner AG, Evans AM, Mito JK, et al. The PANE1 gene encodes a novel human minor histocompatibility antigen that is selectively expressed in B-lymphoid cells and B-CLL. Blood. 2006;107(9):3779–3786.

37. Rijke B de, Horssen-Zoetbrood A van, Beekman JM, et al. A frameshift polymorphism in P2X5 elicits an allogeneic cytotoxic T lymphocyte response associated with remission of chronic myeloid leukemia. J. Clin. Invest. 2005;115(12):3506–3516.

38. Torikai H, Akatsuka Y, Miyazaki M, et al. A Novel HLA-A*3303-Restricted Minor Histocompatibility Antigen Encoded by an Unconventional Open Reading Frame of Human TMSB4Y Gene. J. Immunol. 2004;173(11):7046–7054.

39. Hahn T, Sucheston-Campbell LE, Preus L, et al. Establishment of Definitions and Review Process for Consistent Adjudication of Cause-specific Mortality after Allogeneic Unrelated-donor Hematopoietic Cell Transplantation. Biol. Blood Marrow Transplant. 2015;21(9):1679–1686.

40. Wang J, Clay-Gilmour AI, Karaesmen E, et al. Genome-Wide Association Analyses Identify Variants in IRF4 Associated With Acute Myeloid Leukemia and Myelodysplastic Syndrome Susceptibility. Front. Genet. 2021;12:.

41. Tang H, Hahn T, Karaesmen E, et al. Validation of genetic associations with acute GVHD and nonrelapse mortality in DISCOVeRY-BMT. Blood Adv. 2019;3(15):2337–2341.

42. Karaesmen E, Rizvi AA, Preus LM, et al. Replication and validation of genetic polymorphisms associated with survival after allogeneic blood or marrow transplant. Blood. 2017;130(13):1585–1596.

43. Hahn T, Wang J, Preus LM, et al. Novel genetic variants associated with mortality after unrelated donor allogeneic hematopoietic cell transplantation. eClinicalMedicine. 2021;40:.

44. Zhu Q, Yan L, Liu Q, et al. Exome chip analyses identify genes affecting mortality after HLA-matched unrelated-donor blood and marrow transplantation. Blood. 2018;131(22):2490–2499.

45. Witherspoon DJ, Wooding S, Rogers AR, et al. Genetic Similarities Within and Between Human Populations. Genetics. 2007;176(1):351–359.

46. Baldwin RM, Owzar K, Zembutsu H, et al. A Genome-Wide Association Study Identifies Novel Loci for Paclitaxel-Induced Sensory Peripheral Neuropathy in CALGB 40101. Clin. Cancer Res. Off. J. Am. Assoc. Cancer Res. 2012;18(18):5099–5109.

47. Jurtz V, Paul S, Andreatta M, et al. NetMHCpan-4.0: Improved Peptide–MHC Class I Interaction Predictions Integrating Eluted Ligand and Peptide Binding Affinity Data. J. Immunol. 2017;199(9):3360–3368.

48. Scholtalbers J, Boegel S, Bukur T, et al. TCLP: an online cancer cell line catalogue integrating HLA type, predicted neo-epitopes, virus and gene expression. Genome Med. 2015;7:118.

49. Cornaby C, Montgomery MC, Liu C, Weimer ET. Unique Molecular Identifier-Based High-Resolution HLA Typing and Transcript Quantitation Using Long-Read Sequencing. Front. Genet. 2022;13:.

50. Zhang M, Sukhumalchandra P, Enyenihi AA, et al. A novel HLA-A*0201 restricted peptide derived from cathepsin G is an effective immunotherapeutic target in acute myeloid leukemia. Clin. Cancer Res. Off. J. Am. Assoc. Cancer Res. 2013;19(1):247–257.

51. Terai YL, Huang C, Wang B, et al. Valid-NEO: A Multi-Omics Platform for Neoantigen Detection and Quantification from Limited Clinical Samples. Cancers. 2022;14(5):1243.

52. Choo JAL, Liu J, Toh X, Grotenbreg GM, Ren EC. The Immunodominant Influenza A Virus M158–66 Cytotoxic T Lymphocyte Epitope Exhibits Degenerate Class I Major Histocompatibility Complex Restriction in Humans. J. Virol. 2014;88(18):10613–10623.

53. Pidala J, Kim J, Schell M, et al. Race/ethnicity affects the probability of finding an HLA-A, -B, -C and -DRB1 allele-matched unrelated donor and likelihood of subsequent transplant utilization. Bone Marrow Transplant. 2013;48(3):346–350.

