## Supplemental and Supplemental Figure Legends for "Shared Graft Versus Leukemia Minor Histocompatibility Antigens in DISCOVeRY-BMT"

**Supplemental Methods:**

SNP Typing by PCR Amplification and Sanger Sequencing

PCR primers for the SNP loci encoding the novel mHAs that were validated by mass spectrometry were designed using Primer3 and Geneious. Sequencing primer T7 pro sequence was added to the 5’ end of each forward primer for ease of sequencing. Genomic DNA was extracted from frozen PBMC pellets from alloSCT recipient pre-transplant and post-transplant D90 peripheral blood draw samples from samples expression HLA A*02:01, HLA B*35:01, or HLA C*07:02, obtained from the UNC Lineberger Comprehensive Cancer Center Tissue Procurement Facility, using the Qiagen DNeasy Blood and Tissue Kit (Qiagen, 69504) and purified using the Monarch DNA Cleanup Kit (NEB, T1030S). D90 samples were taken to represent alloSCT donor genetic information, as patient samples were selected for successful engraftment, meaning that at this time point the sample is >95% donor as transplant has engrafted and hematopoiesis is fully donor in origin. PCR amplification was performed using the FastStart High Fidelity PCR System (Roche, 3553400001). Samples were directly sent for Sanger Sequencing by Eton Biosciences using T7pro sequencing primer. Genotyping is shown in Supplemental Figure 5.

**Supplemental Figures:**

Table 1: Patient demographic characteristics of the DISCOVeRY-BMT study. Listed characteristics were selected based on relevance for antigen prediction work and clinical association assessments.

Supplemental Figure 1: HLA typing for AML cell lines. A) shows Venn diagram of number of typed HLA alleles agreed upon and disagreed upon by HLA typing on TRON database versus performed by UNC Hospitals Clinical Laboratories. B) shows alleles called by one source only. C) shows alleles agreed upon by both HLA typing sources.

Supplemental Figure 2: Differential Gene Expression analysis and HLA alleles expressed by cell lines commonly used in presented experiments. Expression levels were calculated relative to the average expression in these five cell lines and top 30 genes that differed from the average were shown. Several cell lines for each of the three studied HLA alleles were examined and one selected based on ease of expansion to needed number of cells for mass spectrometry.

Supplemental Figure 3: Assessment of presence or absence of SNPs encoding validated predicted GvL mHAs. 3a) shows number of copies of alleles encoding fourteen novel HLA A*02:01 mHAs in seven DRPs from alloSCT for AML. Two novel mHAs were excluded due to difficulty of appropriate primer design. Two previously known HLA A*02:01-binding mHAs from the literature, HA-2 and UTA2-1, are also shown. 3b) shows number of copies of alleles encoding three novel HLA B*35:01 mHAs in one DRP. 3c) shows number of copies of alleles encoding five novel HLA C*07:02 mHAs in four DRPs.

Supplemental Figure 4: Gating strategy for mHA immunogenicity assessment. Gating was performed on single cells, lymphocytes, and live cells. A) shows negative control cells stained with tetramer exposed to UV light with no peptide. B) shows positive control cells from CD8 T cells coculture with flu-M1_58-66_ pulsed DCs, stained with flu-M1_58-66_ tetramer. C) shows cells from CD8 T cell coculture with novel mHA UNC-HEXDC-V stained with UNC-HEXDC-V tetramer.
