## Supplemental Figures for "Shared Graft Versus Leukemia Minor Histocompatibility Antigens in DISCOVeRY-BMT"

| **Category** | **Recipient Characteristics** | **First Cohort (n=2357)** | **Second Cohort (n=874)** | **Total (n=3231)** |
|---|---|---|---|---|
| Age, years | <=40<br>>40 | 961 (41)<br>1396 (59) | 322 (37)<br>552 (63) | 1283 (40)<br>1948 (60) |
| Donor age, years range | | 18-61 | 18-60 | 18-61 |
| Sex | M<br>F | 1331 (56)<br>1026 (44) | 480 (55)<br>394 (45) | 1811 (56)<br>1420 (44) |
| Donor sex | M<br>F | 1577 (67)<br>780 (33) | 620 (71)<br>254 (29) | 2197 (68)<br>1034 (32) |
| Disease | AML<br>ALL<br>MDS | 1397 (59)<br>576 (24)<br>384 (16) | 541 (62)<br>121 (14)<br>212 (24) | 1938 (60)<br>697 (22)<br>596 (18) |
| Year of alloSCT | 2000-2002<br>2003-2005<br>2006-2008<br>2009-2011 | 384 (16)<br>848 (36)<br>1125 (48)<br>0 (0) | 33 (4)<br>83 (9)<br>119 (14)<br>639 (73) | 417 (13)<br>931 (29)<br>1244 (39)<br>639 (20) |
| Graft source | Bone marrow<br>Peripheral blood | 1504 (64)<br>853 (36) | 624 (71)<br>250 (29) | 2128 (66)<br>1103 (34) |

A

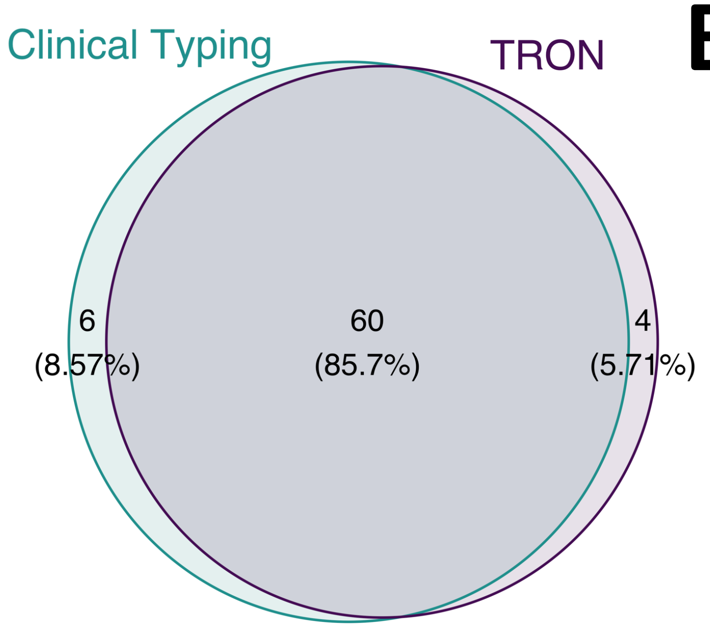


B

| Caller | Cell Line | Allele |
|---|---|---|
| Clinical Typing | OCIM2 | A*01:01 |
| Clinical Typing | U937 | A*31:01 |
| Clinical Typing | Nomo1 | B*55:02 |
| Clinical Typing | OCI-AML3 | B*53:XX |
| Clinical Typing | OCIM1 | C*02:10 |
| Clinical Typing | OCIM2 | C*07:01 |
| TRON | HL60 | A*24:15 |
| TRON | OCI-AML3 | B*53:01 |
| TRON | OCIM1 | C*02:02 |
| TRON | OCIM2 | C*07:06 |

C

| Cell Line | A Alleles | B Alleles | C Alleles |
|---|---|---|---|
| BV173 | 02:01, 30:01 | 15:10, 18:01 | 03:04, 12:03 |
| F36P | 24:02, 33:03 | 40:06, 58:01 | 03:02, 08:01 |
| HL60 | 01:01 | 57:01 | 06:02 |
| MONOMAC1 | 03:01 | 07:02, 51:01 | 07:02, 15:02 |
| NB4 | 11:01 | 35:01, 40:01 | 03:04, 04:01 |
| Nomo1 | 24:02, 26:03 | 54:01 | 01:02, 07:02 |
| OCI-AML2 | 02:01 | 15:01, 18:01 | 03:03, 07:01 |
| OCI-AML3 | 02:01, 23:01 | 44:02 | 04:01, 05:01 |
| OCIM1 | 02:01, 03:01 | 14:02, 15:03 | 08:02 |
| OCIM2 | 02:01 | 08:01, 44:02 | 05:01 |
| U937 | 03:01 | 18:01, 51:01 | 01:02, 07:01 |
| U937A2 | 02:01, 03:01 | 18:01, 51:01 | 01:02, 07:01 |

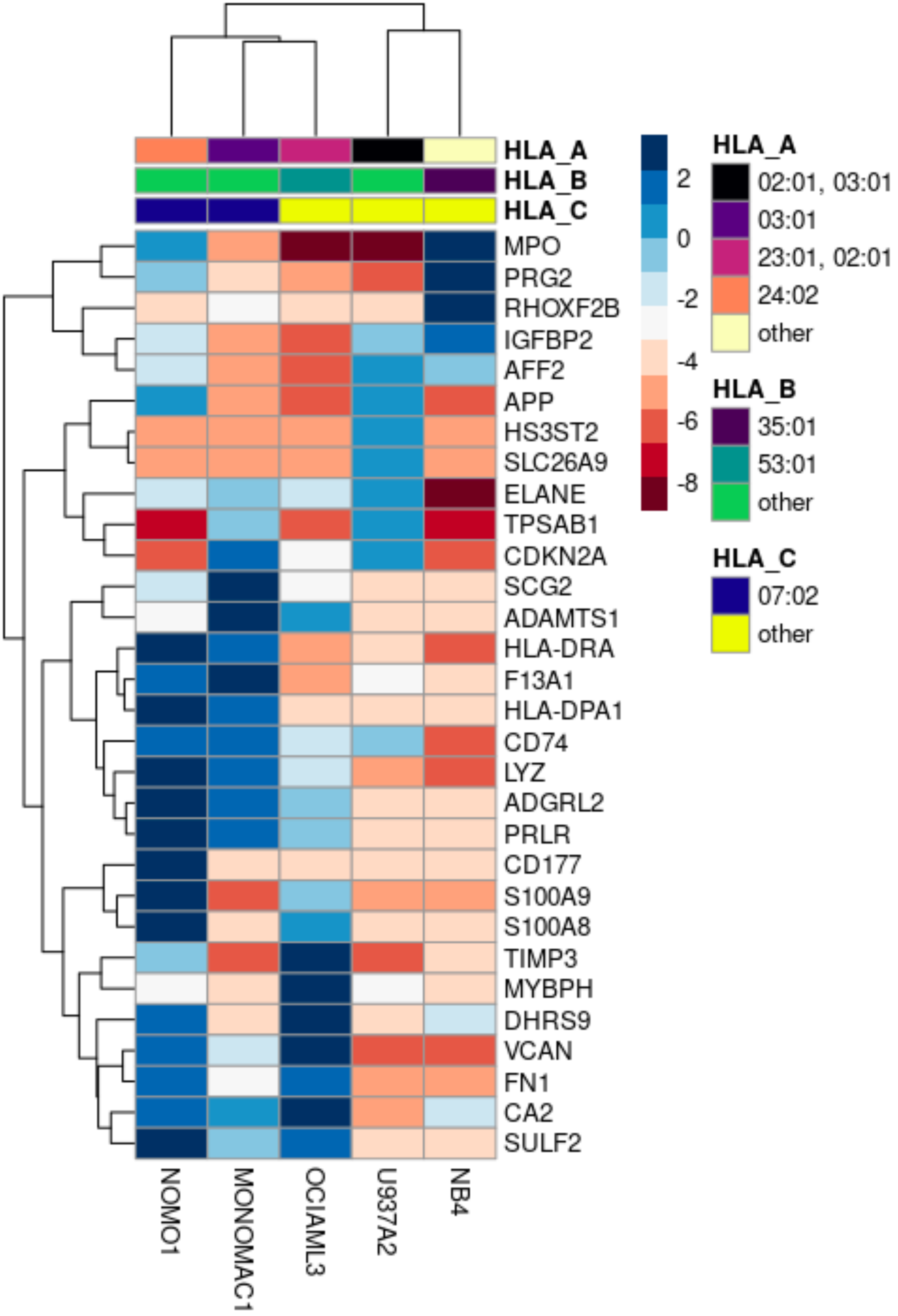
HLA_A
HLA_B
HLA_C
MPO
PRG2
RHOXF2B
IGFBP2
AFF2
APP
HS3ST2
SLC26A9
ELANE
TPSAB1
CDKN2A
SCG2
ADAMTS1
HLA-DRA
F13A1
HLA-DPA1
CD74
LYZ
ADGRL2
PRLR
CD177
S100A9
S100A8
TIMP3
MYBPH
DHRS9
VCAN
FN1
CA2
SULF2
NOMO1
MONOMAC1
OCIAML3
U937A2
NB4
2
0
-2
-4
-6
-8
HLA_A
02:01, 03:01
03:01
23:01, 02:01
24:02
other
HLA_B
35:01
53:01
other
HLA_C
07:02
other

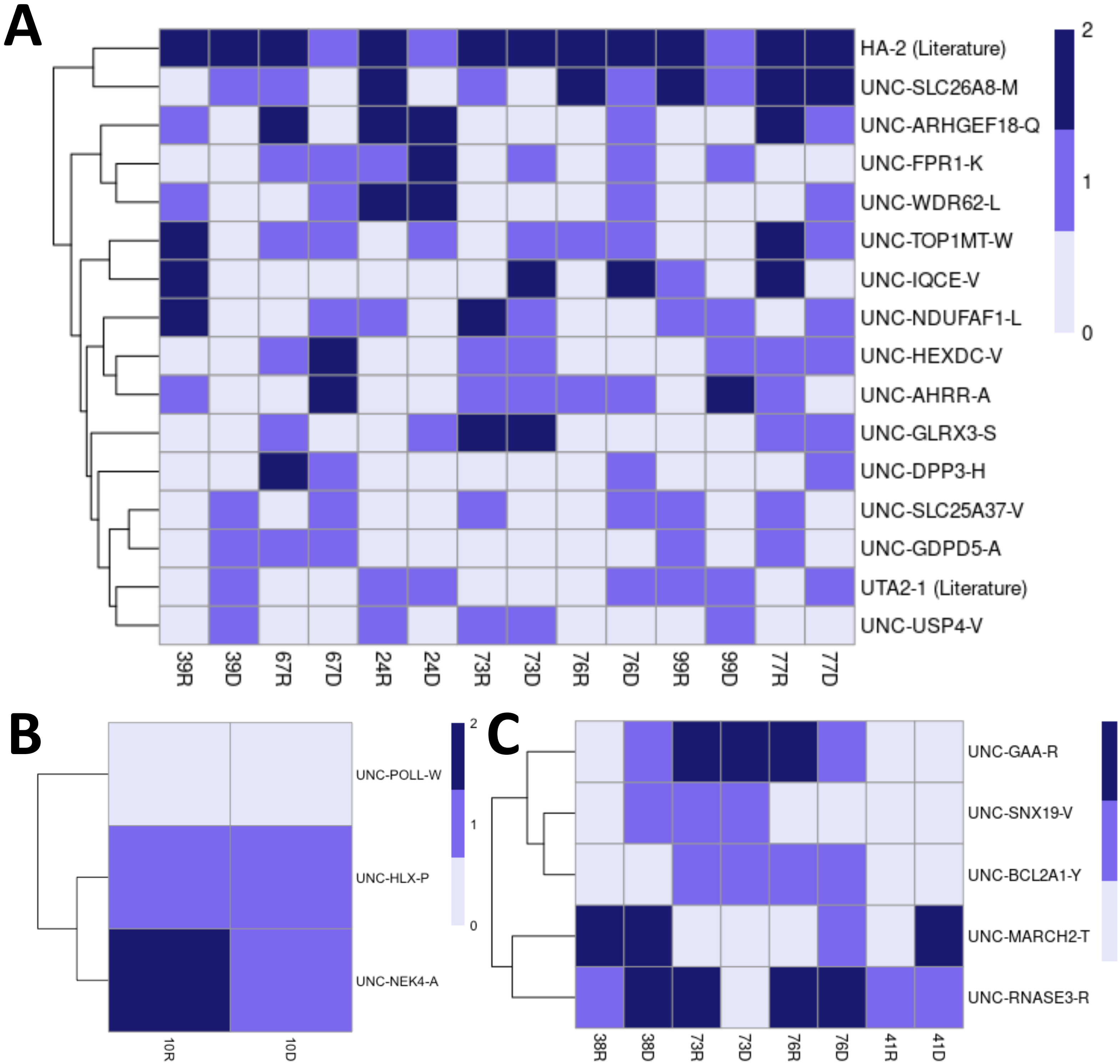
A
HA-2 (Literature)
UNC-SLC26A8-M
UNC-ARHGEF18-Q
UNC-FPR1-K
UNC-WDR62-L
UNC-TOP1MT-W
UNC-IQCE-V
UNC-NDUFAF1-L
UNC-HEXDC-V
UNC-AHRR-A
UNC-GLRX3-S
UNC-DPP3-H
UNC-SLC25A37-V
UNC-GDPD5-A
UTA2-1 (Literature)
UNC-USP4-V
39R
39D
67R
67D
24R
24D
73R
73D
76R
76D
99R
99D
77R
77D
2
1
0
B
UNC-POLL-W
UNC-HLX-P
UNC-NEK4-A
10R
10D
2
1
0
C
UNC-GAA-R
UNC-SNX19-V
UNC-BCL2A1-Y
UNC-MARCH2-T
UNC-RNASE3-R
38R
38D
73R
73D
76R
76D
41R
41D
2
1
0

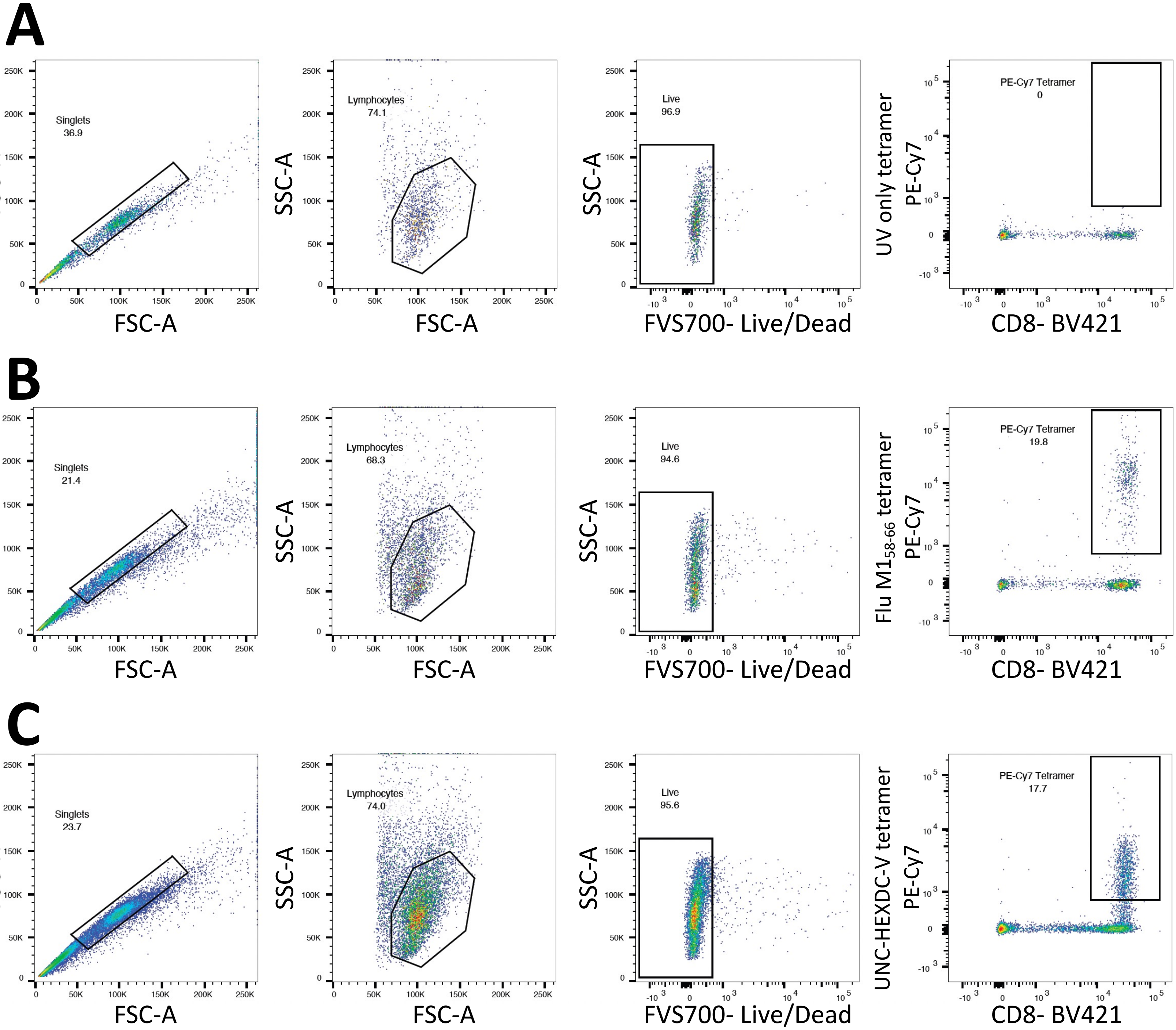
A
Singlets
36.9
FSC-H
FSC-A
Lymphocytes
74.1
SSC-A
FSC-A
Live
96.9
SSC-A
FVS700- Live/Dead
PE-Cy7 Tetramer
0
UV only tetramer
PE-Cy7
CD8- BV421
B
Singlets
21.4
FSC-H
FSC-A
Lymphocytes
68.3
SSC-A
FSC-A
Live
94.6
SSC-A
FVS700- Live/Dead
PE-Cy7 Tetramer
19.8
Flu M1$_{58-66}$ tetramer
PE-Cy7
CD8- BV421
C
Singlets
23.7
FSC-H
FSC-A
Lymphocytes
74.0
SSC-A
FSC-A
Live
95.6
SSC-A
FVS700- Live/Dead
PE-Cy7 Tetramer
17.7
UNC-HEXDC-V tetramer
PE-Cy7
CD8- BV421
